# Whole-body sorcin invalidation does not cause hypothalamic ER stress nor worsens obesity in C57BL/6 male mice fed a westernized diet

**DOI:** 10.1101/2022.09.07.506981

**Authors:** Steven Z. Parks, Guy A. Rutter, Isabelle Leclerc

## Abstract

**Background:** <u>So</u>luble <u>R</u>esistance Related <u>C</u>alcium B<u>in</u>ding Protein (sorcin) is a calcium (Ca^2+^) binding protein which has been shown to play a role in maintaining intracellular endoplasmic reticulum (ER) Ca^2+^ stores and lowering ER stress. Recently, our lab has demonstrated that sorcin expression was downregulated in the islets of Langerhans of mice fed a high-fat diet or in human islets incubated with the saturated fatty acid palmitate. We also showed that overexpression of sorcin under control of the rat insulin promoter (RIP7) in C57BL/6J mice, or whole body sorcin deletion in 129S1/SvImJ mice, improves or impairs insulin secretion and pancreatic β-cell function respectively. The mechanisms behind this beneficial role of sorcin in the pancreatic β-cell might depend on protection against lipotoxic endoplasmic reticulum (ER) stress through improved ER Ca^2+^ dynamics and activation of the Activating Transcription Factor 6 (ATF6) branch of the unfolded protein response (UPR). Whether sorcin is also implicated in hypothalamic ER stress during the progression of obesity is unknown. This could potentially contribute to the diminished satiety typically observed in overweight individuals.

**Aim:** To investigate a potential role of sorcin in hypothalamic ER stress, leptin resistance, hyperphagia and obesity.

**Methods:** Whole-body sorcin null mice, backcrossed onto the C57BL/6J genetic background, were used. Body weight, food intake and EchoMRI body composition were measured in vivo whereas qRT-PCR analysis of sorcin and ER stress markers expression were performed on the arcuate nucleus of the hypothalamus. Leptin signalling through STAT3 phosphorylation was measured by Western blots on sorcin-null HEK293 cells, engineered by CRISPR/Cas9, and transfected with leptin receptor (LepRb).

**Results:** Sorcin expression was not influenced in the arcuate nucleus (ARC) of the hypothalamus by diet-induced obesity. Whole-body sorcin ablation did not cause ARC ER stress nor changes in body weight, body composition or food intake in C57BL/6 male mice exposed to a high-fat, high-sugar diet. STAT3 phosphorylation (Y705) in response to leptin was not impaired in sorcin-null HEK293 cells.

**Conclusion:** In our model, whole body sorcin ablation did not increase hypothalamic ER stress nor influenced food intake or body weight.

## INTRODUCTION

Several studies have shown a relationship between hypothalamic endoplasmic reticulum (ER) stress and leptin resistance, which might be of pathophysiological relevance in the context of obesity (Zhang et al., 2011, Ozcan et al., 2004). Reduction of ER stress with the use of pharmacologic ER inhibitors, such as 4-phenylbutyric acid (4-PBA) and tauroursodeoxycholic acid (TUDCA) have been shown to enhance leptin sensitivity *in-vitro* (Hosoi et al., 2008b), as well as in both diet induced and genetic models of obesity (Çakir et al., 2013).

Sorcin (soluble resistance-related calcium binding protein, gene name SRI, also known as CP-22, CP22, SCN and V19), is a member of the penta-EF-hand family of calcium binding proteins and relocates from the cytoplasm to membranes in response to elevated calcium levels (Colotti et al., 2014, Valdivia, 1998, Ilari et al., 2002, Van der Bliek et al., 1986, Maki et al., 2002, Appelblom et al., 2007, Chen et al., 2018). Sorcin was initially identified in multidrug-resistant cancer cells, is ubiquitously expressed and is highly conserved amongst mammals (Van der Bliek et al., 1986, Meyers and Biedler, 1981) In cardiac myocytes and skeletal muscle cells, sorcin inhibits ryanodine receptor (RyR) activity (Lokuta et al., 1997) and plays a role in terminating Ca^2+^ -induced Ca^2+^ release (Fabiato, 1983, Farrell et al., 2003) an inherently self-sustaining mechanism which, if unchecked, may deplete intracellular Ca_2+_ stores (Stern and Cheng, 2004). Sorcin also activates sarco/endoplasmic reticulum Ca^2+^-ATPase (SERCA) pumps and ensures efficient refilling of ER Ca^2+^ stores after contraction (Matsumoto et al., 2005). In the pancreatic islets of Langerhans and pancreatic beta cells, we first showed that siRNA mediated knockdown of sorcin in the mouse insulinoma MIN6 β-cell line led to a reduction in ER Ca^2+^ stores, showing that the role of sorcin described above is relevant in the context of the pancreatic β-cell (Noordeen et al., 2012). We then showed, both in mouse and human islets, that high-fat diet feeding or incubation with the lipotoxic fatty acid palmitate, causes a reduction in sorcin expression alongside an increase in expression of ER stress markers, whilst overexpression of sorcin was sufficient to reduce ER stress marker expression in palmitate treated islets (Marmugi et al., 2016, Noordeen et al., 2012). We then showed, both in mouse and human islets, that high-fat diet feeding or incubation with the lipotoxic fatty acid palmitate, causes a reduction in sorcin expression alongside an increase in expression of ER stress markers, whilst overexpression of sorcin was sufficient to reduce ER stress marker expression in palmitate treated islets (Marmugi et al., 2016). Moreover, transgenic mice overexpressing sorcin under control of the rat insulin 2 gene promoter (RIP7), expressed in the β-cell, displayed improved glucose tolerance and improved glucose stimulated insulin secretion (GSIS) when fed a high fat diet (HFD). Isolated islets from transgenic mice also display higher glucose induced Ca^2+^ oscillations, and increased ER Ca^2+^ stores, suggesting that improved Ca^2+^ dynamics might be responsible for the *in-vivo* phenotype (Marmugi et al., 2016).

Sorcin is one of the most highly expressed calcium binding proteins in several regions of the brain. Dysregulation of Ca^2+^ signalling is thought to underlie many neurodegenerative diseases, such as Huntingdon’s Disease, Parkinson’s disease and Alzheimer’s disease (Genovese et al., 2020b). Given the role of hypothalamic ER stress in pathologic food intake and leptin resistance and the role of sorcin in maintaining neuronal intracellular Ca^2+^ homeostasis in the prevention of neurodegenerative disorders, it is plausible that sorcin expression might play a role in prevention of lipotoxicity-induced hypothalamic ER stress, leptin resistance and weight gain. Indeed, others also demonstrate a role for sorcin in increased IL-6 stimulated pSTAT3 signalling, albeit in the liver (Li et al., 2017). The JAK2/STAT3 pathway is the canonical pathway of leptin signalling in the hypothalamus, where pSTAT3 binds to the promoters of *pomc* and *agrp,* regulating their transcription. STAT3 increases POMC transcription whilst repressing AgRP; models of neuron specific STAT3 deletion show decreased POMC levels and increased AgRP levels respectively, leading to hyperphagia and obesity (Gao et al., 2004, Li et al., 2017). The JAK2/STAT3 pathway is the canonical pathway of leptin signalling in the hypothalamus, where pSTAT3 binds to the promoters of *pomc* and *agrp,* regulating their transcription. STAT3 increases POMC transcription whilst repressing AgRP; models of neuron specific STAT3 deletion show decreased POMC levels and increased AgRP levels respectively, leading to hyperphagia and obesity (Gao et al., 2004).

Here we wanted to investigate whether sorcin dysregulation may play a role in modulating hypothalamic, and more specifically arcuate nucleus (ARC) ER stress, leading to hyperphagia and weight gain through impaired leptin sensitivity.

## EXPERIMENTAL PROCEDURES

### Animals

All animal procedures undertaken were approved by the British Home Office under the UK Animal (Scientific Procedures) Act 1986 (Project License PPL PA03F7F07 to I.L.) with approval from the local ethical committee (Animal Welfare and Ethics Review Board, AWERB), at the Central Biological Services (CBS) unit at the Hammersmith Campus of Imperial College London.

Animals were housed in a pathogen free facility under controlled temperature (21–23 °C), humidity (45–50%) and 12h light–dark cycle. Unless stated otherwise, mice had free access to normal chow diet and water. Mice were also maintained on alternative diets to induce an obesogenic phenotype; these mice were either fed a high-fat diet (HFD) (Research Diets D12492), consisting of 60% kcal from fat, or a high-fat and high sucrose diet (HFHSD) (Research Diets D12331) consisting of 58% kcal from fat and 25% from carbohydrate sources.

Whole body sorcin-null mice *(Sri^-/-^)* on a 129S1/SvImJ genetic background were generated by homologous recombination as described in (Chen et al.). These mice were backcrossed onto the C57BL/6J background by mating with wild-type C57BL/6J mice (Envigo, Huntingdon, UK) and selecting heterozygous offspring for ≥ 10 generations. Heterozygous *Sri^+/-^* mice were used as breeding pairs to produce animals for our experiments. Genotyping was carried out by PCR as described in (Chen et al.)

### Measurement of Body Weight and Fat Mass

Body weight was recorded weekly and body composition was determined before culling using EchoMRI imaging (EchoMRI-100H Analyzer; Echo Medical Systems LLC, Houston, TX, USA).

### Intraperitoneal Glucose and Insulin Tolerance Tests

Mice were fasted overnight for 16 h for the IPGTT and 8h for the IPITT. Glucose solution (20% D-glucose/water, weight for volume, 1 g/kg body weight) or human regular insulin solution (1 units/kg, catalog no. 19278; Sigma-Aldrich) was administrated intraperitoneally and blood glucose was measured from the tail vein at 0, 15, 30, 60, 90, and 120 min using an ACCU-CHECK Aviva glucometer (Roche).

### Food Intake Studies Post Leptin Administration

Mice were individually housed 24h before the initiation of food intake studies, giving time to acclimatise to their environment. Mice were then fasted for 16h overnight before intraperitoneal administration of recombinant mouse leptin (3μg/g) (R&D System) to half of the cohort and vehicle control (sterile Tris-HCl 20 mM) to the other half. Food was then reintroduced, and cumulative food intake measured for the following 24h, before a washout period of 24h. Experimental conditions were then inverted, with vehicle control mice now receiving leptin administration (3μg/g) and vice-versa, allowing each mouse to act as its own control to assess the effects of leptin on food intake.

### Arcuate Nucleus Microdissection

Mice were euthanized using CO_2_ chambers and death confirmed by cervical dislocation, being careful to avoid damage to the brainstem. Brains were then removed and snap frozen in liquid N_2_ and stored at −80°C. Brains were micro dissected using a previously established protocol, making two coronal incisions, one posterior to the suprachiasmatic nucleus and the other posterior to the hypothalamus, liberating a region approximately 2mm in thickness. The ARC was subsequently dissected from this section using a fine scalpel.

### RNA extraction and qRT-PCR

Arcuate nuclei were then placed in 800μl TRIzol^™^ Reagent (Invitrogen) on ice and homogenized, using Qiagen TissueLyser II at 30Hz for 2 minutes. Homogenized samples were then either used immediately in downstream RNA isolation and processing or stored at −20°C. RNA processing was conducted using TRIzol^™^ Plus RNA Purification Kit according to manufacturer protocol.

RNA solution was then treated with DNase I (RNase-free) (New England Biolabs) to remove genomic DNA from sample and protected against RNA degradation using RNasin^®^ Ribonuclease Inhibitor (Promega). RNA concentration and quality were then measured using a NanoDrop^™^ 2000 spectrophotometer.

Total RNA (500ng to 2μg) was then reverse transcribed using a high-capacity cDNA RT kit (Applied Biosystems) according to the manufacturers’ protocol.

To quantify expression of specific mRNA transcripts, we used SYBR green reagent (Applied Biosystems), and primer pairs to amplify regions corresponding to our genes of interest (Table 1 below). Samples were amplified using Applied Biosystems^™^ 7500 Fast Dx Real-Time PCR Instrument alongside measurement of fluorescent signals. Expression level for each gene was calculated relative to the house-keeping gene *Cyclophilin A*.

**Table 1.**
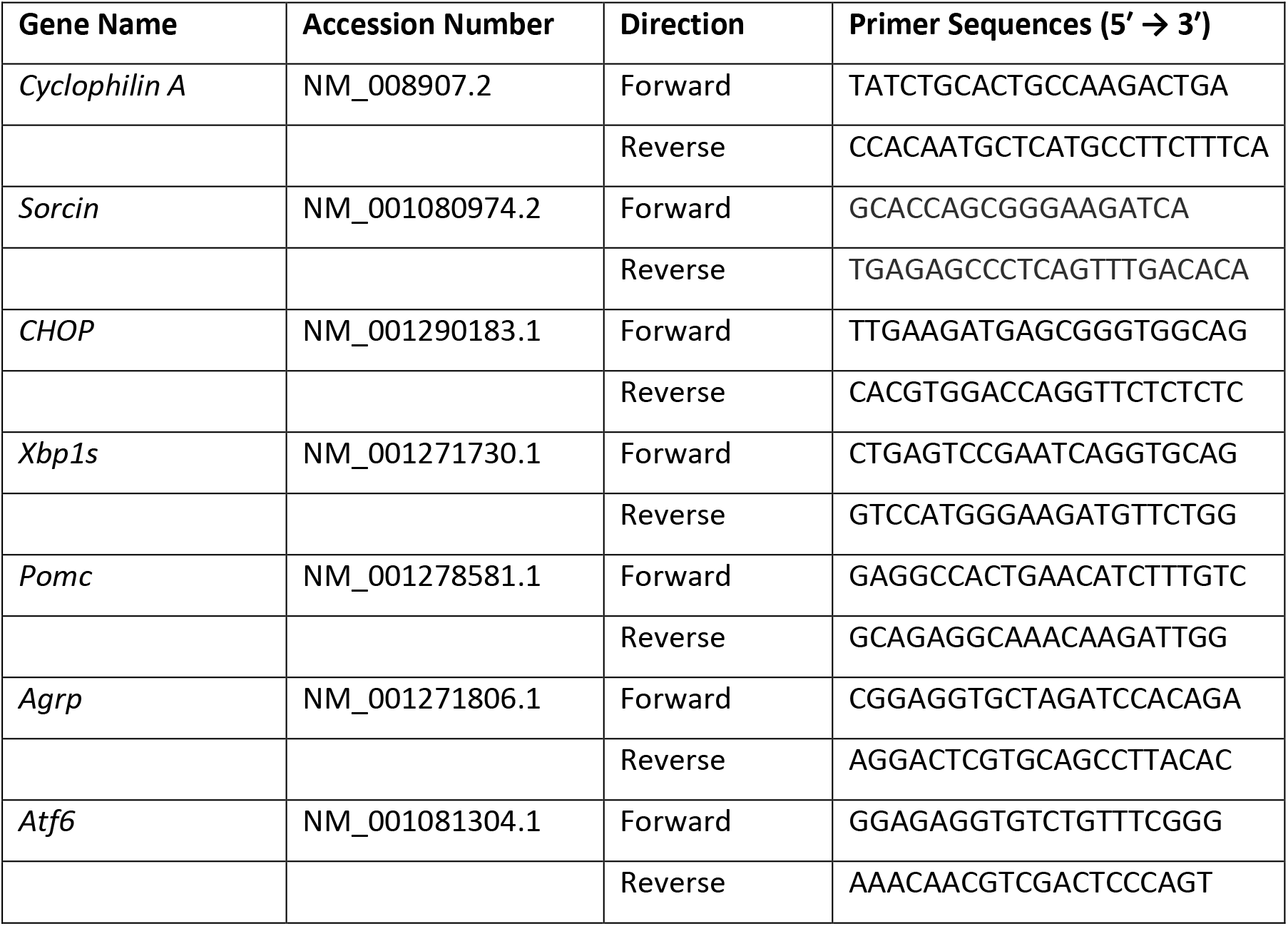
Primers Used for qRT-PCR quantification in *Mus musculus*

**Table 2.** List of antibodies used in Western blots.

| Host species | Antigen | Dilution | Source |
| --- | --- | --- | --- |
| Rabbit | ChREBP | 1:2000 | Abcam NB400-135 |
| Rabbit | Sorcin | 1:5000 | *Described in J Biol Chem.*<br>**278**, 34660-6, 2003 |
| Rabbit | GAPDH | 1:50,000 | Cell Signaling 14C10 |
| Rabbit | Lamin B1 | 1:1000 | Cell Signaling 90875 |
| Rabbit | pSTAT3 (Y705) | 1:5000 | Abcam ab76315 |
| Rabbit | STAT3 | 1:2000 | Abcam ab32500 |

### Mammalian cell culture and transfection

HEK293 (human embryonic kidney) cells (Graham et al., 1977, Rio et al., 1985) were cultured in Dulbecco’s modified eagle medium (DMEM; Sigma Aldrich, Dorset, UK) containing glucose (4.5 g/L), supplemented with penicillin (100 IU/mL), streptomycin (100 μg/mL), L glutamine (4 mM) and FBS (Sigma Aldrich) 10% (vol/vol) in a humidified chamber at 37°C with 5% CO_2_.

Sorcin null HEK293 cells, generated by CRISPR/Cas9 gene editing, have been described in (Parks et al., 2021). HEK293 cells, seeded in 6-well plates, were transfected with 2 μg LepRb-HA plasmid, encoding full length clone DNA of Human leptin receptor, transcript variant 1 with C terminal HA tag (Sino-Biologicals, # HG10322-CY), using Xfect^™^ Transfection Reagent (Takara Bio) according to manufacturer’s instructions. Cells were left for 48h to allow expression of LepRb-HA protein before treatment with recombinant murine leptin (R&D System #498-OB) at 0-10 ng/L for 30 minutes before cell lysis and western blotting for phosphoSTAT3 (Y705), total STAT3 and sorcin as described below.

### Western Blotting

Proteins samples were subjected to sodium dodecyl sulphate–polyacrylamide gel electrophoresis (SDS-PAGE) using 8-10% polyacrylamide gels. Thirty μg of protein sample was run at 120V to separate proteins based on their molecular weights before electrophoretic transfer onto Polyvinylidene Difluoride (PVDF) transfer membranes (Millipore). PVDF membranes containing separated protein samples were blocked for one hour to prevent non-specific primary antibody binding using a solution of 5% skimmed milk, diluted in TBS-Tween (TBST) (50 mM Tris-Cl, 150 mM NaCl, 0.1% vol./vol. Tween 20 (Sigma)). After blocking, membranes were incubated with the primary antibody (see list of antibodies), diluted in 5% skimmed milk TBST at 4°C overnight. The following morning, primary antibody solution was removed, and membrane washed three times in TBST. After washing, membranes were incubated with the horse radish peroxidase (HRP) labelled secondary antibody (see list of antibodies), diluted in 5% skimmed milk TBST at 25°C for one hour. Secondary antibody solution was removed, and membranes washed again three times in TBST. After washing, Clarity ECL Western Blotting Substrate solution (BioRad) was applied to visualise the bands using CL-XPosure^™^ film (Thermo Scientific^™^). Bands were quantified using ImageJ image processing software.

### Statistical Analysis

All data is presented as means ± SEM of at least 3 independent experiments. Statistical significance was analysed by two-tailed unpaired Student’s t-test, one-way ANOVA with Tukey’s multiple comparison test or two-way ANOVA with Šídák’s multiple comparisons test as specified. *, **, *** and **** indicate P < 0.05, 0.01, 0.001 and 0.0001 respectively.

## RESULTS

### HFD Feeding Induces Obesity and Glucose Intolerance in Our DIO Model

To test whether HFD feeding also decreases sorcin expression in the hypothalamus of mice as in their pancreatic islets of Langerhans (Marmugi et al), we used male C57BL/6J mice fed either a control normal chow (NC) diet or a HFD from Research Diets^™^, containing 60% kcal from fat (D12492).

Mice were placed on a NC or HFD at 5 weeks of age and body weight measurements taken regularly to confirm development of obesity in HFD fed mice. In addition to this, body composition was assessed using EchoMRI at the end of the dietary intervention. IPGTTs were conducted at 8, 10 and 16 weeks old to confirm development of glucose intolerance and metabolic dysfunction.

As shown in Figure 1A, mice fed a HFD gained weight significantly more quickly than mice on SD, with significant weight differences developing 5 weeks post HFD introduction (Fig. 1A: 10 weeks of age, SD = 25.78 ± 0.37g vs HFD = 31.25 ± 0.93g, n = 6, p < 0.05, 2-way ANOVA, Šídák’s multiple comparisons test). These differences in weight gain increased with time and at the end of the 13-week study, HFD fed mice on average weighed 12.70g more than SD fed mice (Fig. 1A: 18 weeks of age, SD = 31.13 ± 0.48g vs HFD = 43.83 ± 2.42g, n = 6, p < 0.0001, 2-way ANOVA, Šídák’s multiple comparisons test). Total body fat mass was also measured 13 weeks after diet administration, showing significant differences in body-fat mass between SD and HFD fed mice, suggesting that increased body fat accounts for most of the difference in weight between the two groups. HFD fed mice show 20.03% increase in body-fat percentage compared with SD fed mice, with a body fat percentage of 40.00 ± 2.75 % (Fig. 1B: 18 weeks of age, SD = 19.97 ± 0.57% vs HFD = 40.00 ± 2.75 %, n = 6, p < 0.0001, analysis with unpaired two-tailed t-test).

**Figure 1.**
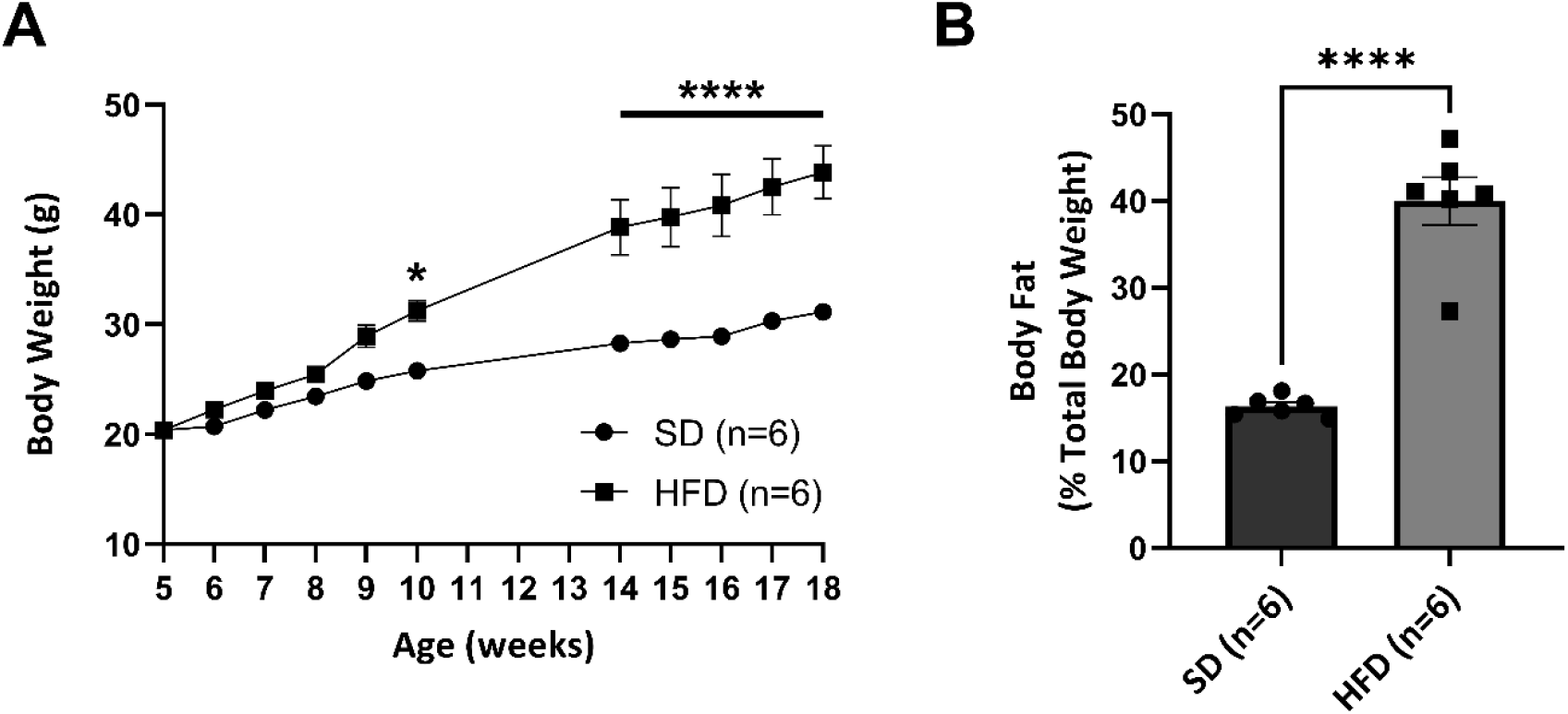
Effect of high-fat diet (HFD) on body weight and fat mass in C57BL/6 wild-type male mice over time. **(A)** Changes in body weight in C57BL/6 wild-type male mice fed a standard diet (SD) or high-fat diet (HFD) over time as indicated (n = 6). **(B)** Body-fat percentage assayed by EchoMRI at 18 weeks old after 13 weeks of dietary intervention (n = 6). Values are means ± SEM. *p < 0.05 and ****p < 0.0001. Data were analysed for significance as follows: A = two-way ANOVA analysis, Šídák’s multiple comparisons test, B = unpaired two-tailed t-test.

As shown in Fig. 2, intraperitoneal glucose tolerance tests (IPGTTs) using 1g glucose/kg body weight were conducted 3 weeks, 5 weeks and 11 weeks post dietary intervention and show a gradual but significant worsening of glucose tolerance throughout this period. HFD fed mice exhibit slightly but significantly altered glucose tolerance at 8 weeks of age, only 3 weeks after dietary intervention when challenged with 1g/kg glucose (Fig. 3A: SD AUC = 1368 ± 52.99 AU vs HFD = 1577 ± 65.44 AU, n = 6, p < 0.05, AUC analysis with unpaired two-tailed t-test). 5 weeks after dietary intervention, HFD fed mice exhibited markedly worsened glucose tolerance, which was exacerbated at 11 weeks post dietary intervention (Fig. 3B: 5 weeks after intervention, SD AUC = 1465 ± 75.71 AU vs HFD AUC = 1800 ± 57.32 AU, n =6, p < 0.01; Fig. 3C: 11 weeks after intervention, SD AUC = 1377 ± 45.00 AU vs HFD AUC = 2079 ± 168 AU, n = 6, p < 0.01, AUC analysis with unpaired two-tailed t-test). Work by the Jackson Laboratory with C57BL/6 mice found comparable body fat percentage and weight at 16 weeks of age, associated with increased triglyceride concentrations, hyperinsulinemia and hyperleptinemia phenocopying early stage Type 2 Diabetes and metabolic syndrome (The Jackson Laboratory, 2022).

**Figure 2.**
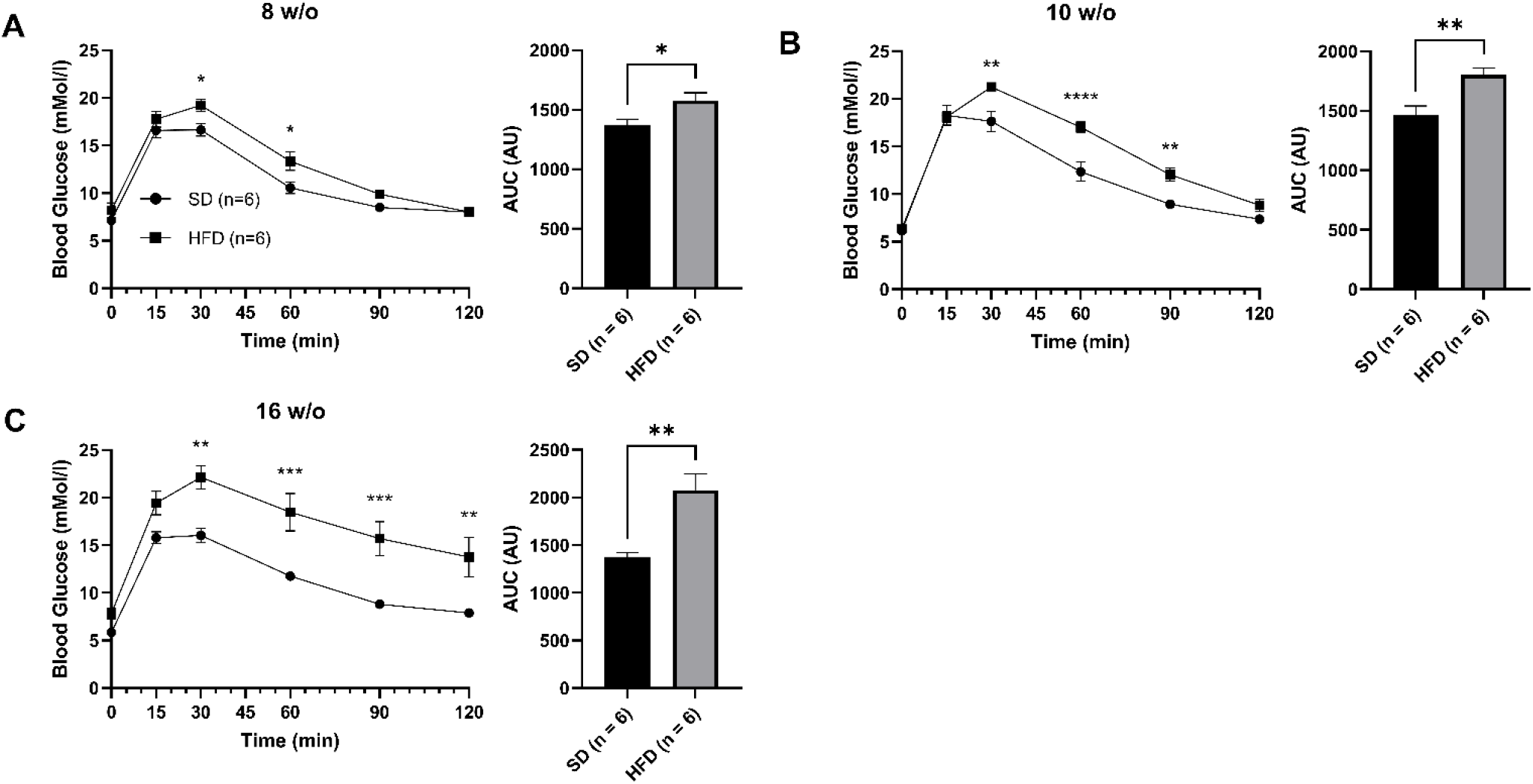
Effect of high-fat diet (HFD) on glucose tolerance in wild-type C57BL/6 male mice over time. **(A-C)** Intraperitoneal glucose tolerance tests (IPGTTs) using 1g/kg glucose were performed on C57BL/6 wild-type male mice on standard diet (SD) or high-fat diet (HFD) at **(A)** 8, **(B)** 10 and **(C)** 16 weeks of age. Area Under Curve (AUC) analyses were conducted and are presented beside. Values are means ± SEM. *P < 0.05; **P < 0.01; ***P < 0.001; ****P < 0.0001. Data were analysed for significance as follows: IPGTTs = two-way ANOVA analysis, Šídák’s multiple comparisons test, AUC = unpaired two-tailed t-test.

**Figure 3.**
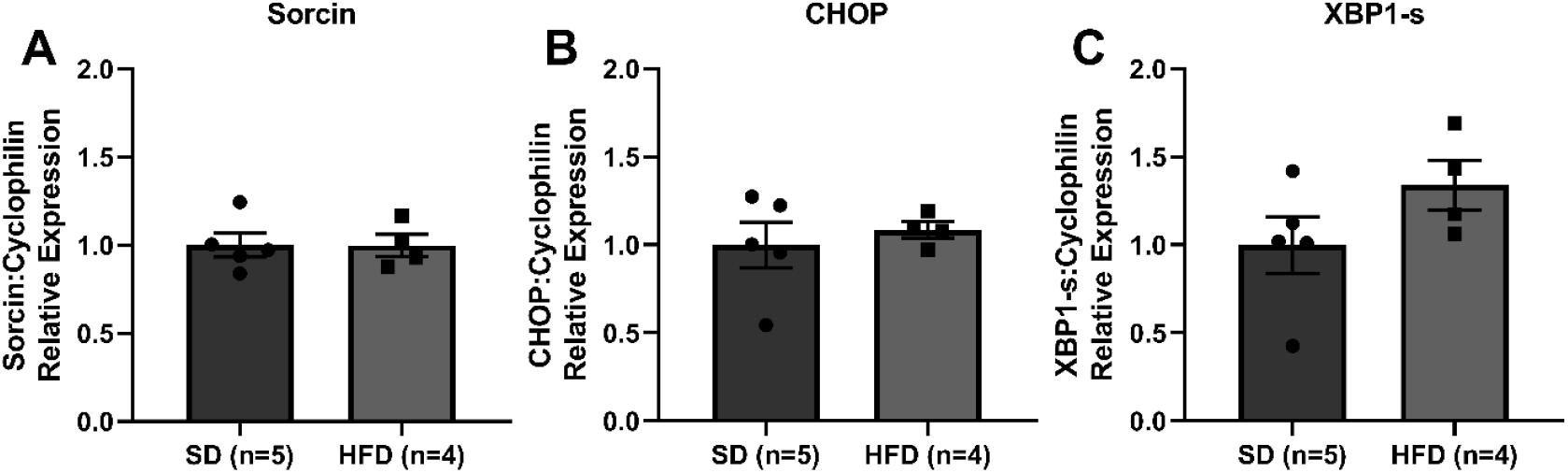
Effect of high-fat diet (HFD) on Sorcin, CHOP and XBP-1 expression in the arcuate nucleus of wild-type C57BL/6 male mice. RT q-PCR analysis shows normalised RNA levels of **(A)** sorcin, alongside ER stress markers **(B)** CHOP and **(C)** XBP1s in arcuate nuclei extracted from standard diet (SD) and high-fat diet (HFD) fed C57BL/6 male wild-type mice, aged 18 weeks of age after 13 weeks of dietary intervention. Values are means ± SEM. Data were analysed for significance using an unpaired two-tailed t-test.

### Sorcin Expression Remains Unchanged in The Arcuate Nucleus of HFD fed Mice

As HFD feeding has been previously linked with hypothalamic ER stress and its association with leptin resistance (Ozcan et al.) we wanted to determine whether reduced expression of sorcin might be responsible for these effects. To determine this, after 13 weeks of dietary intervention, our DIO and control mice were culled and brains were then dissected to isolate the arcuate nucleus (ARC) and RNA extracted for downstream qPCR analysis of gene expression. The results reveal non-significant differences in mRNA levels of sorcin in the ARC (Fig. 3A sorcin: SD = 1.00 ± 0.07 vs HFD = 0.99 ± 0.06, n = 5 vs n = 4, p = 0.98, analysis with unpaired two-tailed t-test). Also, there was no changes in the expression of ER stress marker CHOP and XBP1s in the ARC (Fig. 3B CHOP: SD = 1.00 ± 0.13 vs HFD = 1.09 ± 0.05, n = 5 vs n =4, p = 0.60, analysis with unpaired two-tailed t-test, Fig. 3C XBP1s: SD = 1.00 ± 0.16 vs HFD = 1.34 ± 0.14, n = 5 vs n = 4, p = 0.17, analysis with unpaired two-tailed t-test).

### C57BL/6 Sorcin Null Mice Show Unchanged Body Weight and Fat Mass on HFHSD

Wild-type (*Sri^+/+^*), heterozygous (*Sri^+/-^*) and sorcin null mice (*Sri^-/-^*) on C57BL/6J genetic background were placed on a HFHSD at 6 weeks of age and body weight measurements taken weekly to determine whether *sorcin* null mice showed increased sensitivity to dietary challenge. We found non-significant differences in body weight between across the 12 weeks of HFHSD feeding (Fig. 4 A: 18 weeks of age: *Sri^+/+^* = 38.91 ± 2.29g vs *Sri^+/-^* = 38.66 ± 1.49g vs *Sri^-/-^* = 38.92 ± 1.38g, n = 10 vs 16 vs 12, *Sri^+/+^* vs *Sri^+/-^* p = 0.99, *Sri^+/+^* vs *Sri^-/-^* p = 0.99, *Sri^+/-^* vs *Sri^-/-^* p = 0.99, 2-way ANOVA, Šídák’s multiple comparisons test).

**Figure 4.**
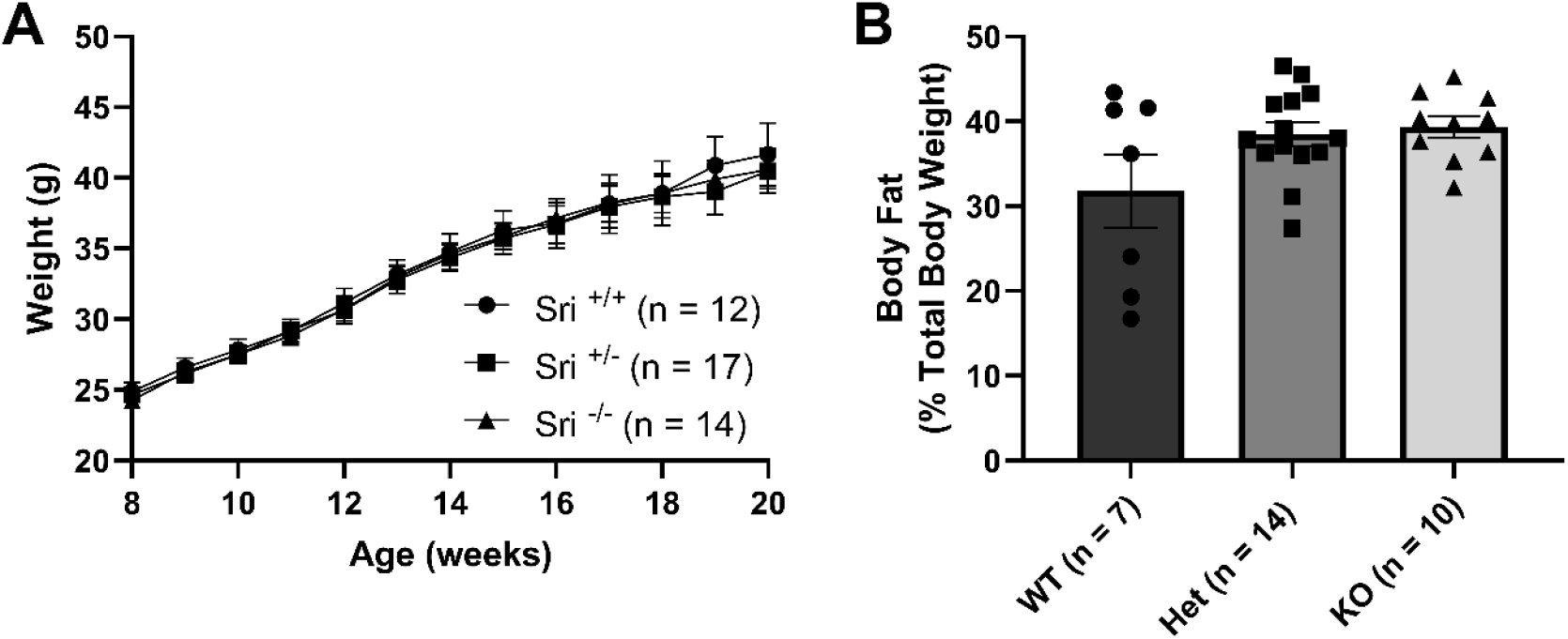
Effect of high-fat high-sugar diet (HFHSD) diet on body weight and fat mass in sorcin WT, Het and KO C57BL/6 male mice. **(A**) Changes in body weight of WT (*Sri^+/+^*), Het (*Sri^+/-^*) and KO (*Sri^-/-^*) sorcin C57BL/6 male mice fed a high-fat high-sugar diet (HFHSD) from 6 weeks of age over time (n = 12 vs 17 vs 14) as indicated. **(B)** Body-fat percentage assayed by EchoMRI at 18 weeks old after 12 weeks of dietary intervention (n = 7 vs 14 vs 10). Values are means ± SEM. Data were analysed for significance as follows: A = two-way ANOVA analysis, Šídák’s multiple comparisons test, B = one-way ANOVA, Tukey’s Multiple Comparison Test.

We also measured body fat percentage at 18 weeks of age using EchoMRI (Figure 4B). These results show a tendency towards increased body fat mass in heterozygous and sorcin null animals, but results are heavily negatively skewed in wild-type mice by three animals and thus show significant deviation (*Sri^+/+^* = 31.81 ± 4.32% vs *Sri^+/-^* = 38.51 ± 1.41g vs *Sri^-/-^* = 39.36 ± 1.27g, n = 7 vs 14 vs 10, *Sri^+/+^* vs *Sri^+/-^* p = 0.10, *Sri^+/+^* vs *Sri^-/-^* p = 0.08, *Sri^+/-^* vs *Sri^-/-^* p = 0.95, 2-way ANOVA, Šídák’s multiple comparisons test).

### C57BL/6 Sorcin Null Mice Exhibit Non-Significant Changes in ARC ER Stress Marker Expression

To determine whether sorcin knockout lead to an increase in ER stress in the ARC nucleus, the main effector of leptin signalling, we measured expression of ER stress markers in RNA isolated from the ARC nucleus of these animals.

As expected, sorcin relative expression was significantly lower in the heterozygous (*Sri^+/-^*) and sorcin null mice (*Sri^-/-^*) compared to wild-type (*Sri^+/+^*) mice (*Sri*^+/+^ = 1.00 ± 0.01 vs *Sri*^+/-^ = 0.66 ± 0.02 vs *Sri^-/-^* = 0.17 ± 0.02, n = 5 vs 12 vs 10, *Sri^+/+^* vs *Sri^+/-^* p < 0.0001, *Sri^+/+^* vs *Sri^-/-^* p < 0.0001, *Sri^+/-^* vs *Sri^-/-^* p < 0.0001, 1-way ANOVA, Tukey’s multiple comparisons test). However, we found non-significant changes in the ER stress markers CHOP, ATF6 and XBP1s in *Sri*^+/-^ and *Sri^-/-^* mice compared to wild-type littermate controls. There was however a tendency towards increased expression of XBP1s suggesting mildly increased ER stress which demands further investigation (*Sri*^+/+^ = 1.00 ± 0.02 vs *Sri*^+/-^ = 1.31 ± 0.11 vs *Sri^-/-^* 1.34 ± 0.12, n = 5 vs 11 vs 9, *Sri^+/+^* vs *Sri^+/-^* p = 0.01, *Sri^+/+^* vs *Sri^-/-^* p = 0.03, *Sri^+/-^* vs *Sri^-/-^* p = 0.96, 1-way ANOVA, Tukey’s multiple comparisons test).

We also investigated changes in expression of the orexigenic and anorexigenic neuropeptide markers AgRP and POMC respectively, to determine if this highlighted any changes in leptin signalling. Our results show non-significant changes in expression of either neuropeptide, consistent with the unchanged body weight and food-intake (Suppl. Fig. 1 POMC: *Sri*^+/+^ = 1.00 ± 0.06 vs *Sri*^+/-^ = 1.20 ± 0.13 vs *Sri^-/-^* 0.84 ± 0.08, n = 6 vs 12 vs 9, *Sri^+/+^* vs *Sri^+/-^* p = 0.47, *Sri^+/+^* vs *Sri^-/-^*p = 0.63, *Sri^+/-^* vs *Sri^-/-^* p = 0.06; AgRP: *Sri^+/+^* = 1.00 ± 0.06 vs *Sri*^+/-^ = 1.20 ± 0.13 vs *Sri^-/-^* 0.84 ± 0.08, n = 6 vs 12 vs 9, *Sri^+/+^* vs *Sri^+/-^* p = 0.47, *Sri^+/+^* vs *Sri^-/-^* p = 0.63, *Sri^+/-^* vs *Sri^-/-^* p = 0.06, 1-way ANOVA, Tukey’s multiple comparisons test)

### C57BL/6 Sorcin Null Mice Show a small increase in food Intake after leptin administration

Given the role of hypothalamic ER stress in leptin resistance, we also wanted to determine changes in leptin sensitivity in *Sri^-/-^* mice. We attempted to do this using pSTAT3 staining of perfusion-fixed coronal brain slices after leptin administration but were unable to optimise the protocol in our mice. Therefore, as an indirect measure of leptin sensitivity, we measured both ad-libitum food intake and food intake after leptin administration in our cohort of mice, on standard chow diet in 8 w/o mice (Fig 6 A, B) or after feeding a HFHSD in 20 w/o mice for 12 weeks (Fig. 6 C, D). We observed non-significant differences at all time-points between *Sri^+/+^, Sri^+/-^* and *Sri^-/-^* mice in ad-libitum food intake at both 8 and 20 weeks of age. Food intake after leptin administration (3 μg/g) also showed no significant difference at most time-points analysed; cumulative food intake at 8 hours after leptin administration was slightly increased in *Sri^-/-^* mice at 8 weeks of age (Fig. 6 B: *Sri^+/+^* = 2.37 ± 0.35g vs *Sri^+/-^* = 2.90 ± 0.29g vs *Sri^-/-^* = 3.08 ± 0.28g, n = 3 vs 9 vs 6, *Sri^+/+^* vs *Sri^+/-^* p = 0.10, *Sri^+/+^* vs *Sri^-/-^* p = 0.03, *Sri^+/-^* vs *Sri^-/-^* p = 0.65, 2-way ANOVA, Šídák’s multiple comparisons test), but this observation disappeared at the 24-hour time point.

**Figure 5.**
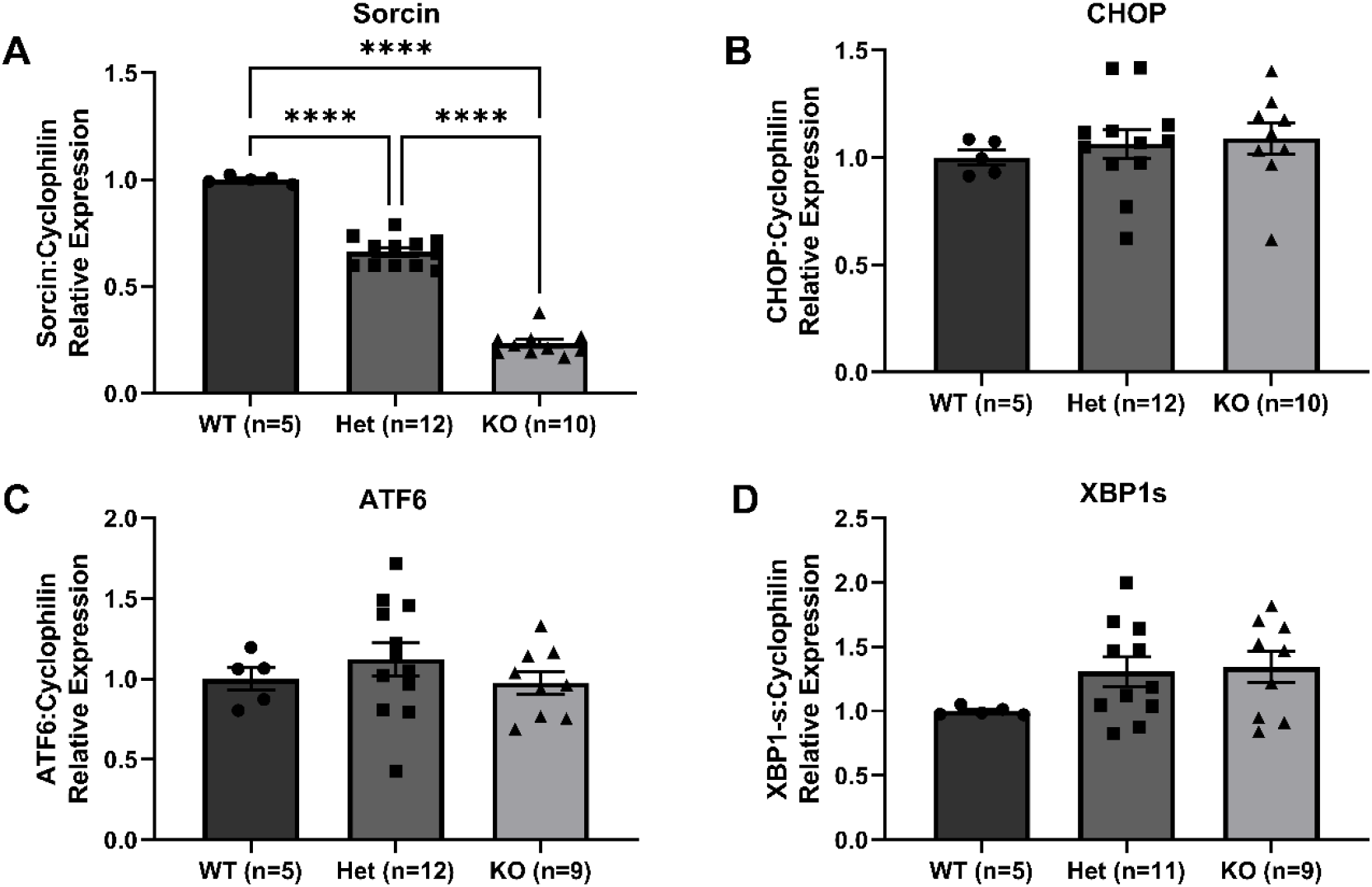
Effect of high-fat high-sugar diet (HFHSD) diet on Sorcin and ER stress markers CHOP, ATF6 and XBP1s expression in the arcuate nucleus of wild-type, heterozygous and null sorcin male mice. RT q-PCR analysis shows RNA expression levels of **(A)** Sorcin, **(B)** CHOP, **(C)** ATF6 and **(D)** XBP1s in the ARC extracted from WT (*Sri^+/+^*), Het (*Sri^+/-^*) and KO (*Sri^-/-^*) C57BL/6 male littermate mice aged 22 weeks old, fed a high-fat high-sugar diet (HFHSD) from age 6 weeks of age. Values are means ± SEM. ****P < 0.0001. Data were analysed for significance using 1-way ANOVA analysis with Tukey’s multiple comparisons test.

**Figure 6.**
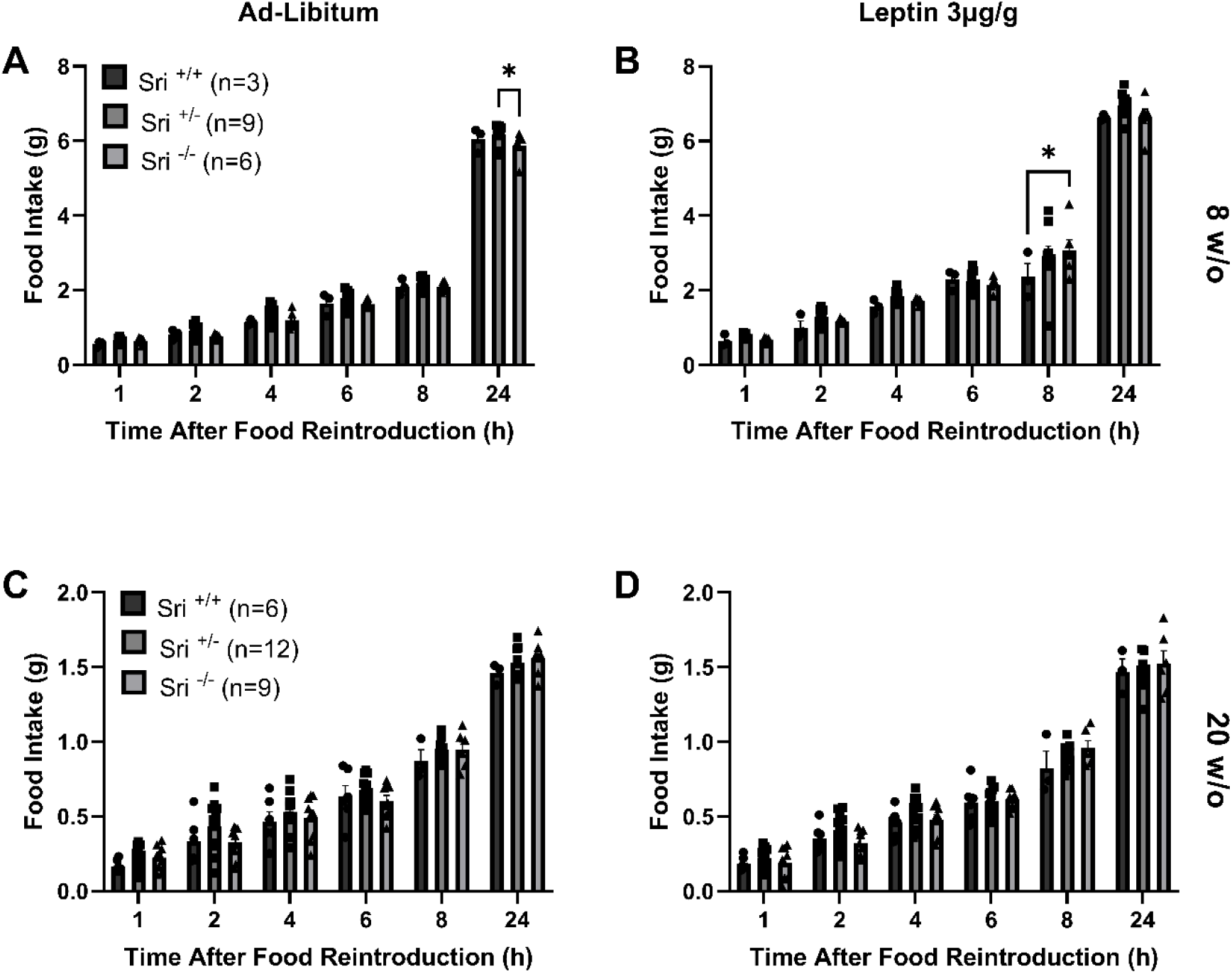
Food Intake Ad-libitum or Post Leptin Administration in sorcin WT, Het and KO C57BL/6 male mice on HFHSD. **(A + C)** Ad-Libitum food intake of littermate C57BL/6 male mice before fed a high-fat high-sugar diet (HFHSD) at **(A)** 8 and **(C)** 20 weeks of age. **(B + D)** Food intake after intraperitoneal leptin administration (3μg/g) of C57BL/6 mice fed a high-fat high-sugar diet (HFHSD) at **(B)** 8 and **(D)** 20 weeks of age. Values are means ±SEM. *P < 0.05. Data were analysed for significance as follows: A = two-way ANOVA analysis, Šídák’s multiple comparisons test, B = one-way ANOVA, Tukey’s Multiple Comparison Test.

### Sorcin Null HEK293-LepRb Cells Show Unchanged Leptin Sensitivity

As previously mentioned, other studies have measured the effects of sorcin depletion or overexpression on STAT3 signalling using *in-vitro* models. Li et al. found that sorcin overexpression in the hepatocellular carcinoma cell line HepG2 increases STAT3 phosphorylation in response to IL-6 stimulation (Li et al., 2017). To determine whether similar mechanisms might occur in response to leptin stimulation, we used our HEK293 sorcin null cells expressing the long-isoform of the leptin receptor (LepRb) by transient transfection with LepRb-HA plasmid. These cells were then stimulated with leptin (0-10 ng/mL), and STAT3 phosphorylation (Tyr705) measured by western blotting of cell lysates. HEK293 wild-type cells and non-transfected cells were used to control against sorcin and LepRb independent STAT3 signalling.

As shown in Fig. 7, there was no significant difference in pSTAT3/STAT3 ratio between wild-type and sorcin null HEK293 cells for any concentration of leptin stimulation. Rather than reduced leptin signalling in sorcin null cells, there is a tendency for increased leptin signalling, with sorcin null cells showing consistently higher pSTAT3/STAT3 ratios for all concentrations of leptin stimulation. Despite this, two-way ANOVA analysis shows non-significant differences between genotypes analysed across all data (wild-type vs Sri null p = 0.15).

**Figure 7.**
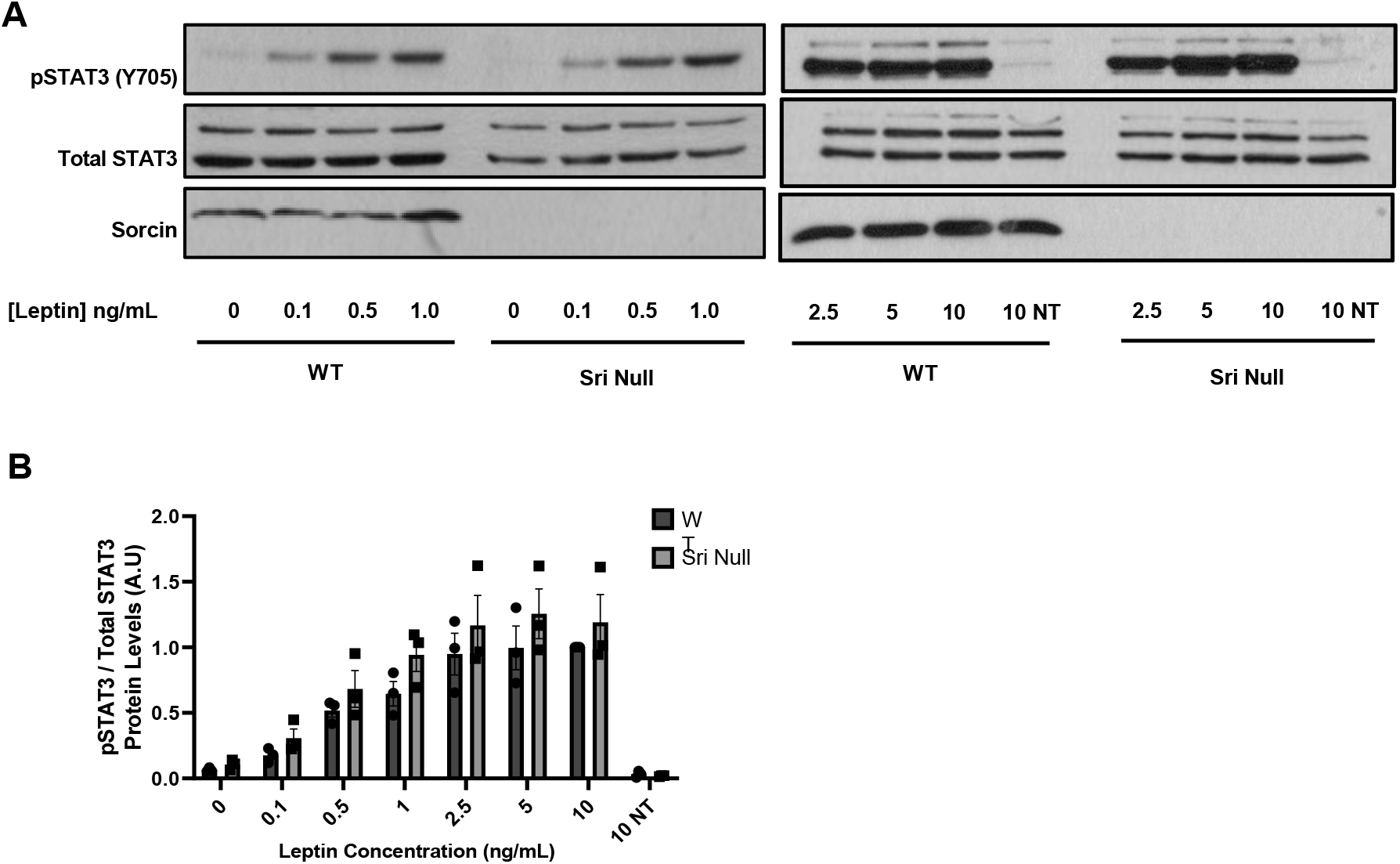
Effect of leptin dose response on STAT3 phosphorylation in LepRb expressing in WT and sorcin-null HEK293 Cells. **(A)** Representative Western (Immuno)blotting of pSTAT3 (Y705), STAT3 and Sorcin from whole protein lysate isolated from wild-type and sorcin null HEK293 cells expressing LepRb and stimulated with leptin (0-1 ng/mL) as described in methods. **(B)** Quantification of Western (Immuno)blotting from 3 independent experiments, showing the ratio between pSTAT3/STAT3 for cells stimulated with leptin (0-10 ng/mL); NT = non-transfected cells. Values are means ± SEM. Data were analysed for significance using 1-way ANOVA analysis with Tukey’s multiple comparisons test.

## DISCUSSION

The role of hypothalamic ER stress in metabolic dysfunction has been the subject of significant investigation, with recent studies demonstrating the relationship between hypothalamic ER stress and leptin resistance, resulting in hyperphagia, positive energy balance and exacerbating metabolic syndrome (Ozcan et al., 2004, Ozcan et al., 2009, Çakir et al., 2013, Zhang et al., 2008, Hosoi et al., 2008b). Reduction of hypothalamic ER stress in mouse models of obesity, using ER stress inhibitors such as 4-PBA or TUDCA, has been shown to rescue leptin sensitivity and highlights possible therapeutic avenues. However, the underlying cause of ER stress induction in this context has not yet been identified. Since we previously showed that sorcin expression was reduced in pancreatic islets of Langerhans and HEK293 cells in response to palmitate exposure, and that sorcin protected against ER stress (Marmugi et al., Parks et al.), we hypothesised that altered expression of sorcin, as one of the most highly expressed calcium binding proteins in the brain, in the context of HFD feeding, might be sufficient to lead to hypothalamic ER stress and altered leptin signalling. Additionally, sorcin has been shown to interact with STAT3 in the liver, increasing its phosphorylation, providing an alternative mechanism by which sorcin might improve leptin signalling in the hypothalamus.

However, we observed no effect of HFD on sorcin expression within the arcuate nucleus of the hypothalamus (Fig. 3) despite significant increases in body weight, fat mass and glucose intolerance (Fig. 1, 2). These results suggest that changes in hypothalamic sorcin expression are not responsible for changes in leptin sensitivity associated with obesity in our murine model. However, as the changes in ARC expression of ER stress markers CHOP and XBP1s after 12 weeks of HFD feeding was insignificant, these results do not exclude the role of sorcin in inhibiting ARC ER stress.

We further investigated the role of sorcin in modulating energy homeostasis. *Sri^-/-^* mice were backcrossed onto the C57BL/6J strain to investigate whether sorcin plays a role in preventing leptin resistance using a well-characterized model of diet-induced obesity. Wild-type (*Sri^+/+^*), heterozygous (*Sri^+/-^*) and sorcin null (*Sri^-/-^*) littermate male mice were administered a HFHSD from 8 weeks of age to identify any changes in whole-body energy homeostasis. We observed non-significant alterations in body weight, fat mass or food intake in these animals despite robust decreases in sorcin expression in the heterozygous (*Sri^+/-^*) and sorcin null (*Sri^-/-^*) animals (Fig. 4, 5, 6). Our LepRb expressing HEK293 sorcin null *in-vitro* model corroborated these observations, showing non-significant differences in leptin-induced STAT3 signalling (Fig. 6).

In conclusion, we were not able to show a regulation of sorcin expression in the arcuate nucleus of the hypothalamus in HFD fed mice and sorcin null mice did not show any changes in whole-body weight, global food intake or hypothalamic mRNA levels of ER stress markers compared to their heterozygous or wild-type littermates. Moreover, STAT3 phosphorylation in response to leptin signalling was not affected by the absence of sorcin in HEK293 cells.

---

**Suppl. Figure 1.**
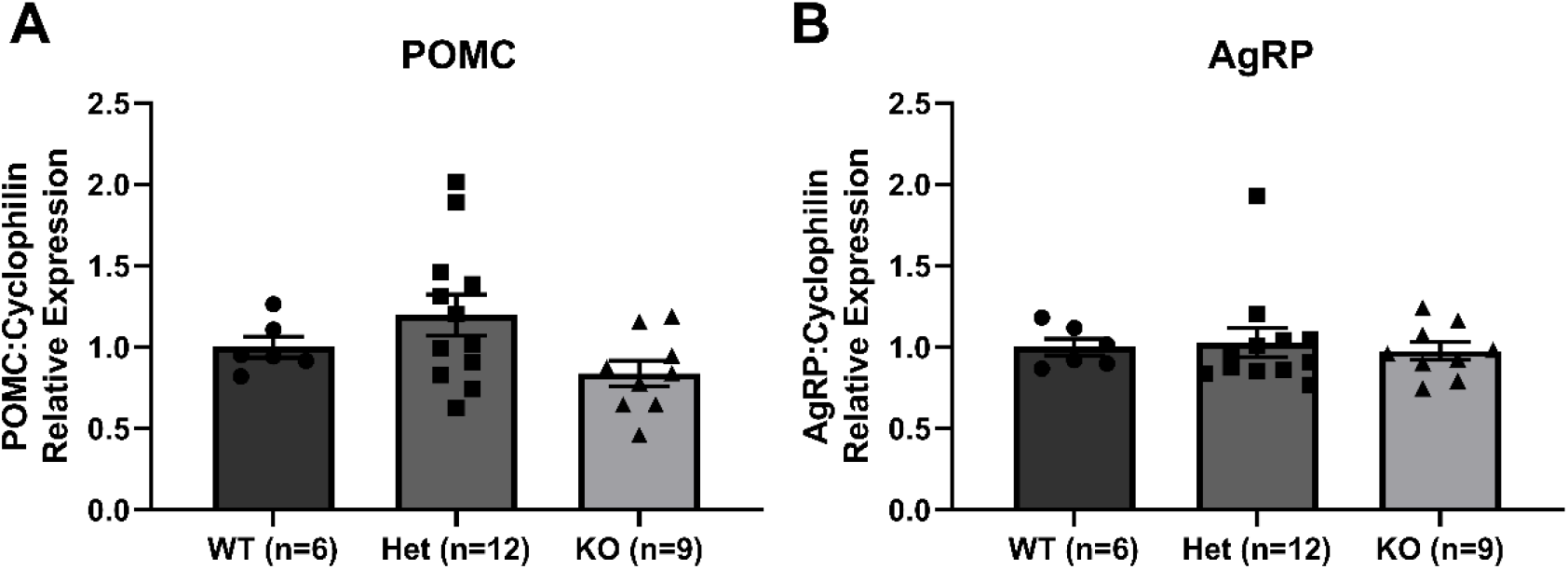
Sorcin Null Mice Have Unchanged Expression of Orexigenic/Anorexigenic Signals AgRP and POMC. RT q-PCR analysis shows RNA expression levels of **(A)** POMC and **(B)** AgRP in the ARC extracted from *Sri^+/+^, Sri^+/-^* and *Sri^-/-^* C57BL/6 mice fed a high-fat high-sugar diet (HFHSD). Values are means ± SEM. Data were analysed for significance using 1-way ANOVA analysis with Tukey’s multiple comparisons test.

## Authors’ contributions

SZP and IL designed, performed experiments and analysed data. GAR and IL contributed resources. SZP and IL wrote the manuscript. IL is the guarantor of this work and takes responsibility for the integrity and accuracy of the data presented.

## Acknowledgements and Funding

Funding was provided by Imperial College London to SZP (President’s PhD studentship) and grants to IL from Diabetes UK (BDA:12/0004535 and BDA:16/0005485) and The Rosetrees Trust (A1063); to GAR from the Wellcome Trust (WT098424AIA and 212625/Z/18/Z), the MRC (MR/R022259/1, MR/J0003042/1, MR/L020149/1 and MR/N00275X/1) and Diabetes UK (BDA/15/0005275 and BDA 16/0005485).

